# Neural correlates of experience with CCTV surveillance of naturalistic prosocial and antisocial interactions: a reverse correlation analysis

**DOI:** 10.1101/691790

**Authors:** Julia A. Gillard, Karin Petrini, Katie Noble, Jesus A. Rodriguez Perez, Frank E. Pollick

## Abstract

Previous research using reverse correlation to explore the relationship between brain activity and presented image information found that Face Fusiform Area (FFA) activity could be related to the appearance of faces during free viewing of the Hollywood movie “The Good, the Bad, and the Ugly” (Hasson, et al, 2004). We applied this approach to the naturalistic viewing of unedited footage of city-centre closed-circuit television (CCTV) surveillance. Two 300 second videos were used, one containing prosocial activities and the other antisocial activities. Brain activity revealed through fMRI as well as eye movements were recorded while fifteen expert CCTV operators with a minimum of 6 months experience of CCTV surveillance alongside an age and gender matched control group of fifteen novice viewers were scanned while watching the videos. Independent scans functionally localized FFA and posterior Superior Temporal Sulcus (pSTS) activity using faces/houses and intact/scrambled point-light biological motion displays respectively. Reverse correlation revealed peaks in FFA and pSTS brain activity corresponding to the expert and novice eye movements directed towards faces and biological motion across both videos. In contrast, troughs in activation corresponded to camera-induced motion when a clear view of visual targets were temporarily not available. Our findings, validated by the eye movement data, indicate that the predicted modulation of brain activity occurs as a result of salient features of faces and biological motion embedded within the naturalistic stimuli. The examination of expertise revealed that in both pSTS and FFA the novices had significantly more activated timeframes than the experienced observers for the prosocial video. However, no difference was found for the antisocial video. The modulation of brain activity, as well as the effect of expertise gives a novel insight into the underlying visual processes in an applied real-life task.

## Introduction

Human action understanding as a hallmark of social interaction encompasses the active monitoring and decoding of human body movement in everyday dynamic environments in order to draw inferences about people’s current state of mind, mood or intent (Grafton and Tipper, 2012). Decoding and recognizing potential threats, such as violent intent or hostile behaviour in context-dependent social interactions is especially important in scenarios where normal boundaries of behaviour are more likely to be crossed (Eldridge and Roberts, 2008). For this reason, Close Circuit Television (CCTV) cameras and operators are tasked with monitoring everyday scenes and unfolding interactions to ensure the safety of the community and the individual (Webster, 2009). In the UK, there is one CCTV camera for every 32 people, from street corners and alleyways to shop fronts and high streets, covering both public and private premises. With incoming information from an estimated five million machines across the UK in 2004 alone, there is an oversaturation of footage or CCTV data being recorded on a 24/7 basis. This positions the UK as a leading user of CCTV for safety and crime investigation purposes (Norris et al., 2004).

As a result, local councils and private security firms require CCTV operators for common surveillance tasks such as the monitoring and controlling of incidents (Keval and Sasse, 2008). The accurate detection and identification of violent or hostile intent and operation of CCTV cameras is performed simultaneously to liaising with security and emergency services. However, with a ratio of CCTV operators to camera monitors varying from 16:100 to 3:90 (Keval and Sasse, 2006), the online perceptual load is ostensibly high given multiple dynamic scenes and a limited range of sensory data (Tickner and Poulton, 1975). As a consequence, behavioural and fMRI studies have been conducted to investigate the effects of expertise on viewing and evaluating CCTV footage. Evidence suggests that experienced operators, compared to novices use a more systematic pattern of eye movements (Roffo et al., 2013) when scanning scenes and exhibit higher sensitivity in judgments of hostile intent and suspiciousness (Petrini et al., 2014). These behavioural differences are in conjunction with differences in brain activity as revealed by fMRI data using a whole brain analysis of experienced operators and novices when they viewed 16 second clips of different intentions. Results of this study showed decreased activity in the anterior parahippocampal gyrus of operators suggesting more efficient neural processing in judging hostile intentions (Petrini et al., 2014).

Although previous behavioural studies have used CCTV footage to examine the recognition of intention (Howard et al., 2011; Koller et al., 2015; Troscianko et al., 2004), previous work aimed at establishing the functional neural correlates of perceiving these types of social signals have largely used perceptually constrained stimuli, such as isolated presentations of static facial images or biological motion represented by brief point light displays (Johansson, 1973). While such studies have revealed a widespread action observation network involving temporal, parietal and frontal brain areas (Gardner et al., 2015; Saygin, 2004), two areas have received much attention for their role in the early processing of faces and bodies. Face processing reliably activates a region of fusiform cortex known as the face fusiform area (FFA) (Berman et al., 2010; Kanwisher and Yovel, 2006), while the processing of biological motion is thought to activate areas within the posterior superior temporal sulcus (pSTS) (Allison et al., 2000; Grossman et al., 2000; Grossman and Blake, 2002). However, there is limited evidence of whether these findings generalise to complex, dynamic social scenes extended over long time periods.

The use of experimentally controlled and un-naturalistic stimuli does not necessarily reflect the complexity of real-world dynamic naturalistic scenes underlying social judgments. As a result, researchers have argued that insights into the functionality of individual brain areas relating to face or body motion processing should rather be ascertained from realistic visual stimuli under free viewing conditions compared to constrained and directed viewing of isolated stimuli (Hasson et al., 2010, 2004). This approach has been implemented in a venture of neuroscience (Hasson et al., 2004) and cinematic research (Smith, 2006). Hollywood movies represent complex dynamic stimuli that can be easily presented to individuals while recording both brain activity and eye fixations under free viewing conditions. As part of this approach, Hasson et al. (Hasson et al., 2004) used reverse correlation to explore the neural similarity as a function of attentional synchronicity between viewers’ gaze. Reverse correlation allows for particular features of a naturalistic scene to be associated with common activations in brain activity across all subjects, revealing the synchronicity of brain activity in relationship to function. From this work, Hasson et al. (Hasson et al., 2004) were able to show that areas such as the face fusiform area (FFA) respond to the unconstrained viewing of faces within a scene, even when the facial stimuli are embedded in complex and dynamic scenes. However, Hollywood movies are nonetheless subject to careful editing procedures that direct attention of the viewer towards the narrative of the story (Smith et al., 2012). In other words, close up shots of faces and interactions implicitly direct the viewer’s attention away from the contextual features and onto the relevant stimuli in a similar manner as experimentally constrained visual stimuli (Smith, 2006).

This study aims at further exploring the neural underpinnings of facial and biological motion processing in uncontrolled naturalistic stimuli. Using close-circuit television (CCTV) footage, which has been gathered from different cities across the UK, the stimuli represent a naturalistic scenario of nighttime activities and interactions between single individuals and small groups in major urban centres. Exploring brain activity of areas relevant to social processing while viewing the complexity of embedded features within a dynamic scene allows for a naturalistic inspection of brain activity as opposed to controlled presentation of visual stimuli in an established hypothesis driven analysis in which pre-defined stimuli are implemented to activate local brain areas. In contrast, the novel analysis methods of reverse correlation and intersubject correlation posited by Hasson et al. (Hasson et al., 2004) allows for an unbiased analysis of localized brain activity, as activation patterns draw attention to specific features within stimuli. A limitation of this analysis in establishing the functionality of brain regions has been the association of video edited close-up shots of faces or objects with peaks in activation.

In this research we use a two-step process to first functionally localise the FFA and pSTS regions and then to examine the timecourse of activity in these regions for novice and experienced CCTV operators when they view prosocial and antisocial videos. By examining FFA we will see if we can replicate the findings of Hasson et al. (Hasson et al., 2004) using our dynamic CCTV footage. The objective of this study is to gain a greater insight into the relationship and diverse functionality of biological motion processing and action understanding under naturalistic conditions of prosocial and antisocial interactions. Additionally, we aim to extend our knowledge of the function of pSTS and FFA by examining the modulatory effect of experience with CCTV footage. Most of what is known about the neural mechanisms of biological motion perception has been derived from typical observers in experimentally constrained paradigms, and by studying experienced observers of naturalistic interactions we can gain insight into whether these mechanisms generalise to typical observers. The hypothesis in the first instance is that reverse correlation will reveal peaks in FFA activation upon viewing faces and facial features, while dips in FFA activation will correspond to an absence of eye fixations on faces or facial features or the presence of camera movements. In a similar line of thought, a second hypothesis predicts that reverse correlation will also reveal peaks and dips in pSTS activation that correspond to the perceptual availability and fixation on biological motion or human movement. Furthermore, the peaks in pSTS activation will be selective to biological motion as opposed to non-biological motion. This is relevant as the CCTV footage used includes camera actions related to the original camera operator performing actions such as zooming in and out to gather evidence and switches of camera as the observed individuals move through the cityscape. Finally, based on previous findings [11] and the CCTV operators’ familiarity with this type of material, we predict reduced peaks of activation in CCTV operators when compared to novice viewers, and different responses in these groups to pro- and anti-social movies.

## Materials and Methods

### Participants

Fifteen expert CCTV operators (4 female, age range 38-57, mean age = 47.13, SD = 7.01) with a minimum of 6 months experience of CCTV surveillance (range 0.8-17 years, mean experience = 10.35, SD = 5.33) participated in this study alongside an age and gender matched control group of fifteen novice viewers (4 female, age range 38-59 years, mean age = 47.73, SD = 6.32). There was no significant difference between the age of the expert and novice viewers (independent t-test, t(28) = 0.246, p = 0.807). The novices had no prior experience of using CCTV surveillance to monitor human behaviour in their work environment. Informed consent was obtained and ethical approval granted by the UK Ministry of Defence Research Ethics Committee (Petrini et al., 2014).

### Design

The study was a within- and between-subject mixed design. The independent factors were group expertise (expert and novice viewers), as well as movie type (antisocial and prosocial). The dependent measures were the brain activity as revealed by fMRI while viewing the movies as well as eye movement data acquired during scanning.

### Stimuli

CCTV footage of an evening urban environment was obtained from CCTV control centres throughout the UK and examined to produce two 300-second movies with either anti-social or pro-social interactions between a set of people. This footage was similar to that used by Petrini et al. (Petrini et al., 2014) that restricted the segment duration to a series of 16 second movies, with each segment judged to be anti- or prosocial by a group of naïve participants.

The CCTV footage termed the “antisocial” movie starts by focusing on a group of four young males, of which two appear to be having a verbally aggressive argument which escalates into physically aggression while the two other associates attempt to intervene to deescalate the hostile situation between the confronting individuals. As the group progress through the inner city, the CCTV camera follows the scene from an empty street to more populated streets and finally to a largely crowded space with several individuals and groups. The four individuals proceed through the crowd when policemen on foot and horseback apprehend the two individuals involved in the violent incident.

In the other 300-second CCTV footage termed “prosocial” movie, the camera follows a small group of young men conversing in a relaxed manner as they are walking along empty pavements, until reaching a larger open public space. They start saying goodbye, hugging and engaging in joking gestures and casual conversation for several minutes until finally separating and walking in smaller groups into different directions. The camera continues to follow three individual members of the group as they leave the scene along another street.

To facilitate understanding what events might give rise to the measured brain activity both movies were annotated to find the start and end of various kinds of events. These included events common to both movies such as camera motion and unique events like fight (antisocial) and joking (prosocial). This annotation was performed by the first author (JG) prior to fMRI data analysis based on event categories derived from predominant behavioural actions by the protagonists apparent from visual inspection.

### Neuroimaging and eye tracking procedure

After providing informed consent, participants underwent a series of scans, including the passive watching of the antisocial and prosocial CCTV movies discussed here. Scanning was performed in two sessions on a 3T Siemens (Erlangen, Germany) Tim Trio MRI scanner equipped with a 12-channel head coil. Whole-brain echo-planar functional images were acquired while participants passively watched, through a mirror, the movies back-projected to a screen behind the scanner. The movies were presented sequentially with their order counterbalanced among observers. Data for each movie was acquired in separate scans separated by a short break of approximately one minute. The functional scans consisted of one run for each movie (TR = 2000 ms; TE = 30 ms; 32 Slices; 3 mm3 isovoxel; 70X70 image resolution; 150 Volumes), plus the functional localiser runs (two runs for each localiser) to identify the fusiform face area (FFA) and the biological motion processing area posterior Superior Temporal Sulcus (pSTS). The details of the functional localiser runs are identical to those described in (McAleer et al., 2014). Functional data consisted of high-resolution T1 weighted anatomical scans using MPRAGE sequences (192 slices; 1mm^3^ Sagittal Slice; TR = 1900 ms; TE = 2.52 ms; 256 × 256 image resolution).

For the prosocial and antisocial displays, participants were instructed to passively view the movies as they appeared on screen. The displays were displayed on a Windows PC with stimulus presentation being controlled using the software package Presentation, Neurobehavioral Systems (www.neurobs.com). The CCTV footage had been converted to an AVI format with 480 × 576 pixel resolution with minimal loss of quality. The movies were presented to participants centred in the screen at a visual angle of 30°×22.5°, while the movies occupied a visual angle of approx. 19°×22.5°. Each movie presentation was preceded and followed by a blank screen followed by a 10 second fixation towards a central target. While watching the CCTV movies, the position of the right eye was recorded at a sampling rate of 60 Hz for each participant, in line with recommendations when using the *ViewPoint Eye Tracker*® *PC 60* by Arrington Research, Inc. (www.ArringtonResearch.com).

### Functional MRI Data Analysis

Analysis of the brain imaging data was performed using BrainVoyager QX (Brain Innovation B.V., Maastricht, Netherlands). After standard pre-processing of the brain images including Slice Scan Time Correction; 3D Motion Correction; and Temporal Filtering; the individual brains were aligned into stereotaxic space [22].

#### Localiser Scans

The two functional localizer scans were analysed for each individual participant to identify a region in the posterior Superior Temporal Sulcus (pSTS) sensitive to biological motion (Beauchamp et al., 2002; Grossman et al., 2000; Grossman and Blake, 2002; Zacks et al., 2006) as well as a face sensitive region in the fusiform gyrus (Kanwisher et al., 1997; McCarthy et al., 1997) known as the fusiform face area (FFA). The pSTS functional localiser included visual conditions of a static point light displays, intact biological motion displays and scrambled biological motion displays. Similarly, the FFA functional localiser included the visual conditions of visual noise, face images and house images [21].

#### Reverse Correlation Analysis

Following localisation of the FFA and pSTS areas we analysed the timecourse of brain activity for these two regions to see how it varied with experience and movie type. The time courses of localized brain activation for each hemisphere were extracted from the peak voxels within the regional clusters of activation in the pSTS and FFA areas obtained. The time courses were averaged across hemispheres, z-normalized, and smoothed using a three point moving average (Hasson et al., 2004). The hemodynamic response delay can potentially impede the accurate interpretation of the blood-oxygen-level-dependent (BOLD) signal and analysis of brain activity using reverse correlation in particular during the presentation of fast paced movie sequences (Hasson et al., 2004). To account for this, participant’s time courses were shifted by a hemodynamic delay of six seconds to better approximate the movie presentation (Hasson et al., 2004).

Independent sample t-tests were performed for each time point along the averaged time course across all participants’ individual time series. Time points significantly different from the average mean value (p<0.05) reflect times when there was consistent brain activation across viewers and these time points were considered further in the reverse correlation analysis. In the reverse correlation analysis we compared the nature and frequency of the movie frames which evoked significant peak activation and deactivation between expert and novice viewers, and between the antisocial and prosocial movies for both FFA and pSTS brain activity. The aim of this study was to associate visually salient features with localized peak brain activation unbiased by pre-existing notions of functional properties. To facilitate this aim, the significantly correlated peaks and dips in activation were matched to the event and the eye position movement data used to verify the gaze location at particular instances. For example, if at the beginning of an argument there was a peak in FFA activation then it could be checked if participants were looking at the face at this point in time.

### Eye Movement Analysis

Eye movement data was collected to allow for the qualitative validation of significant peaks and troughs of brain activity in the reverse correlation analysis. Unfortunately, due to technical difficulties the extent of eye movement data recorded was limited to only a subset of 10 observers (5 novices and 5 experienced participants) who had a complete data set of eye movements for both movies. Scan paths were created for each individual and superimposed on the original movies, in which the sequence of the viewer’s eye fixations was visualized within each timeframe of the movie. Due to the dynamic nature of the stimuli, the scanpaths obtained from the eye tracking procedure allowed for a real-time investigation of the direction of gaze and attention for each viewer throughout the antisocial and prosocial movie. The direction and spatial localization of viewer’s fixation was used to associate visually salient features with brain activity in the pSTS and FFA.

## Results

### Localisation of FFA and pSTS

The localiser scan worked in the majority of participants to specify the FFA and pSTS brain regions. The average talairach coordinates and the number of participants were localisation was successful are presented in Table 1.

**Table 1.** Average Talairach Coordinates for fMRI Localiser Brain Activation in mm (SD)

| Localiser | Group | Right Hemisphere |  |  |  |  |  |  | Left Hemisphere |  |  |  |  |  |  |
| --- | --- | --- | --- | --- | --- | --- | --- | --- | --- | --- | --- | --- | --- | --- | --- |
|  |  | N | x |  | y |  | z |  | N | x |  | y |  | z |  |
| STS | Novices | 10 | 50.3 | (8.1) | -43.4 | (12.3) | 9.9 | (4.6) | 8 | -48.3 | (6.1) | -54.5 | (10.6) | 14.1 | (5.1) |
|  | Experts | 8 | 50.6 | (6.4) | -38.5 | (9.3) | 8.9 | (8.4) | 11 | -52.3 | (5.9) | -41.6 | (14.9) | 9.7 | (5.7) |
| FFA | Novices | 10 | 35.0 | (5.3) | -50.0 | (15.8) | -15.3 | (3) | 8 | -39.3 | (7.2) | -45.9 | (3.9) | -17.3 | (3.5) |
|  | Experts | 7 | 37.9 | (3.8) | -45.3 | (7.3) | -17.1 | (3.8) | 8 | -39.8 | (4) | -48.9 | (3.2) | -14.9 | (3.3) |

### Event Annotation

The relationship between event categories of each movie and brain activity in pSTS and FFA is illustrated in Fig 1 and Fig 2. As can be seen, the perceptual availability of faces and/or dynamic movements was modulated by the differential pace and content of the movie, reflected in particular in the use of cinematic devices, such as large-scale camera movements. These cinematic devices affected both movies equally, whereas the intensity and pace of social interactions differed between movies.

**Fig 1.**
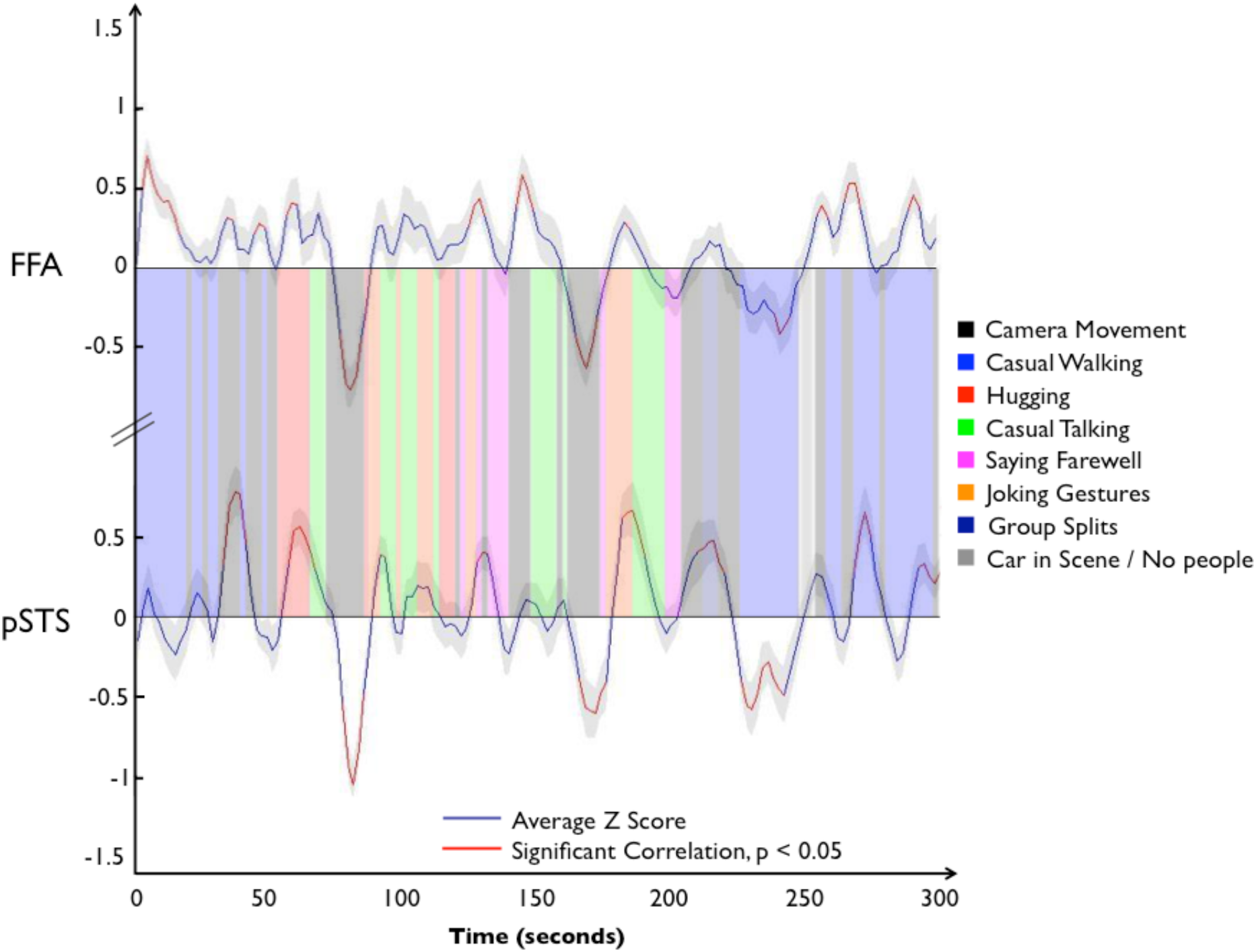
Prosocial Event Categories in Relation to FFA and pSTS Activity. The average FFA and pSTS activity of all viewers, as well as annotated events categories were overlaid on to the prosocial time course to illustrate the associations between peaks and troughs in activation and corresponding events categories. Peaks and troughs in brain activity significantly different from the mean value across participants (p<0.05) are marked in red.

**Fig 2.**
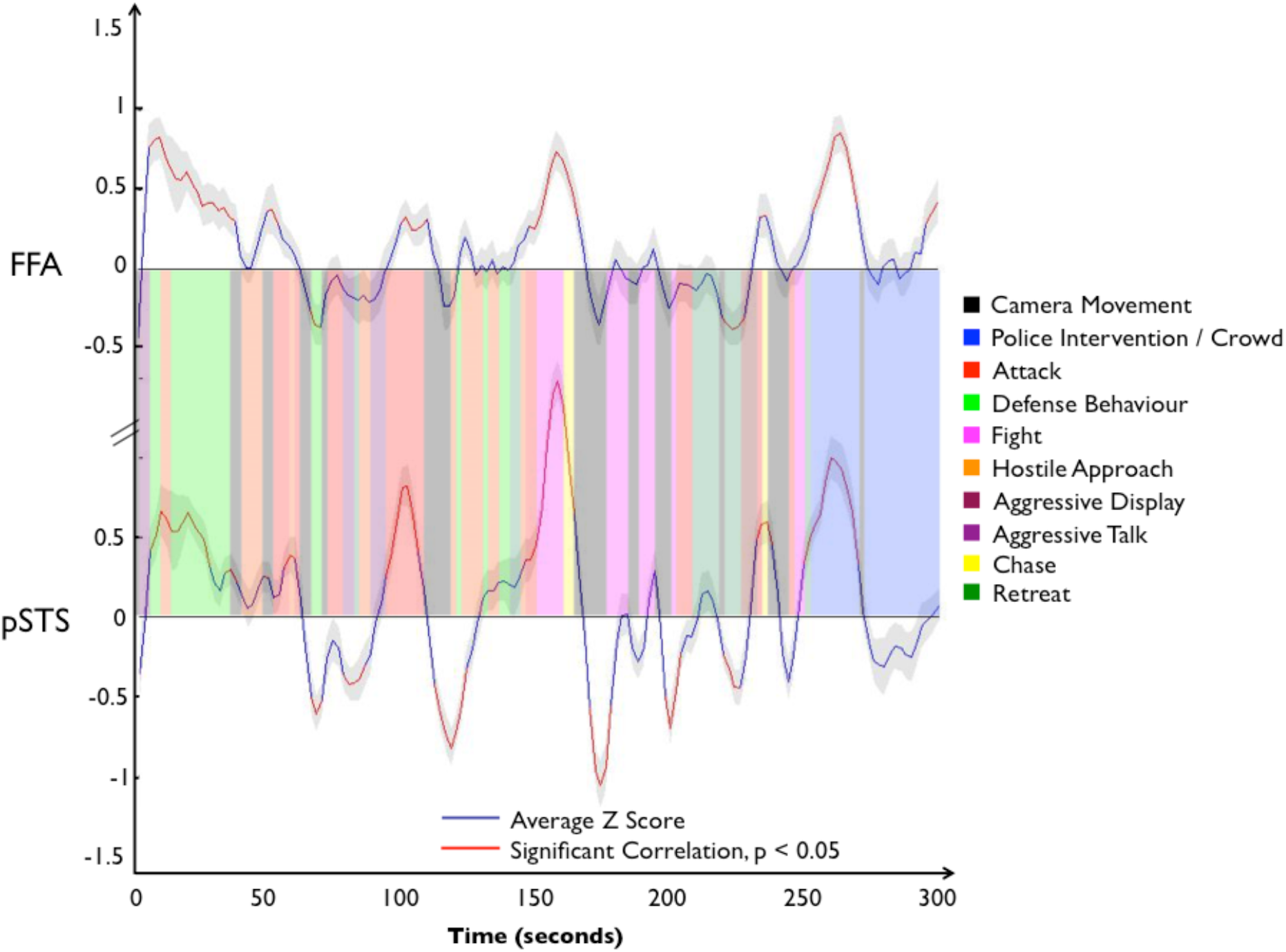
Antisocial Event Categories in Relation to FFA and pSTS Activity. The average FFA and pSTS activity of all viewers, as well as annotated events categories were overlaid on to the antisocial time course to illustrate the associations between peaks and troughs in activation and corresponding events categories. Peaks and troughs in brain activity significantly different from the mean value across participants (p<0.05) are marked in red.

### Reverse Correlation Analysis

Despite large sections of the movie being dominated by camera movements, particular features of scenes relating to face or motion processing validated through viewer’s fixation points were found to be closely associated with the peaks and dips in localized brain activations in pSTS and FFA. The time-series of pSTS and FFA activation for the prosocial and antisocial movie are illustrated in Fig 3 including the times and peak values that were statistically significant.

**Fig 3.**
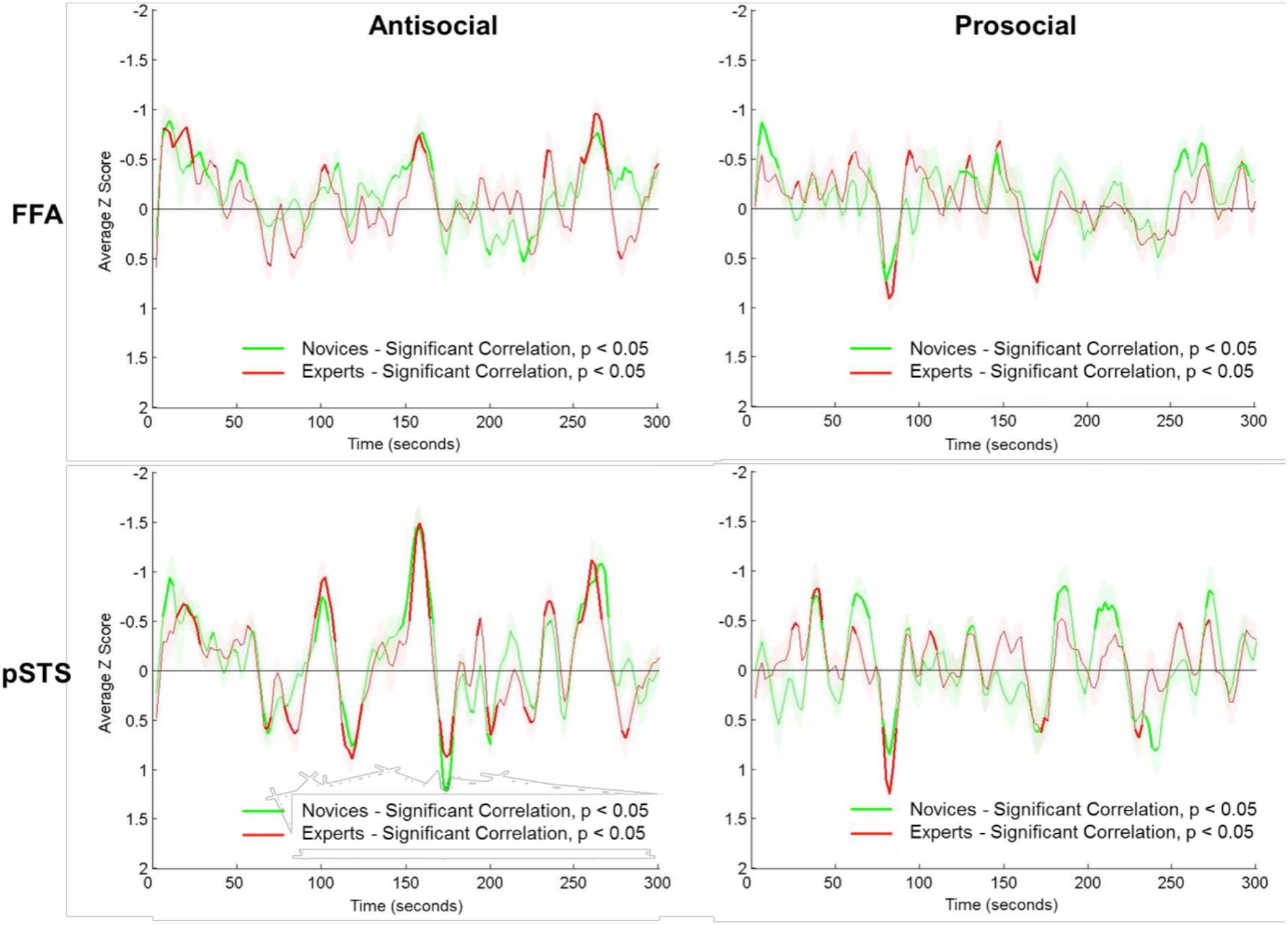
Reverse Correlation in FFA and pSTS for Novices and Experts. Peaks and troughs in brain activity significantly different from the mean value across participants (p<0.05) in FFA and pSTS for the prosocial and antisocial movie are illustrated in bold for expert viewers in red and novice viewers in green, respectively.

#### Face Fusiform Area (FFA)

The relationship between significant peaks in FFA activation and the visual perception of faces in experts and novices viewing the prosocial movie and antisocial is illustrated in

Fig 4 and Fig 5 respectively. The red line denotes the significant peak activity at p<0.05, the blue line denotes the average Z score across viewers. Eye fixations of expert viewers are denoted by individually coloured dots in still images derived from the CCTV footage. Occasional eye movements were missing from the footage if eye fixations temporarily strayed outside of the image or eye gaze was occluded due to blinking. The still images presented here have been edited to remove identifying information, while preserving general information of the scene. The still images correspond to indicated time points reflecting increased or decreased level of activation. Large camera movements and/or the limited perceptual availability of faces or facial features resulted in dips in activation as illustrated by a series of timeframes depicting zooming and panning camera movements. In contrast, eye fixations centred on the presence of faces within the scene are reflected in significant peaks in FFA activation. As opposed to predominantly large sweeping camera movements within the prosocial movie, the antisocial movie is characterised by more frequent, rapid camera adjustments following the aggressive interactions of the two main protagonists. This is reflected in less severe decreases in activations during the prosocial movie in contrast to more frequent decreases in activation during the antisocial movie across both groups (See Fig 3).

**Fig 4.**
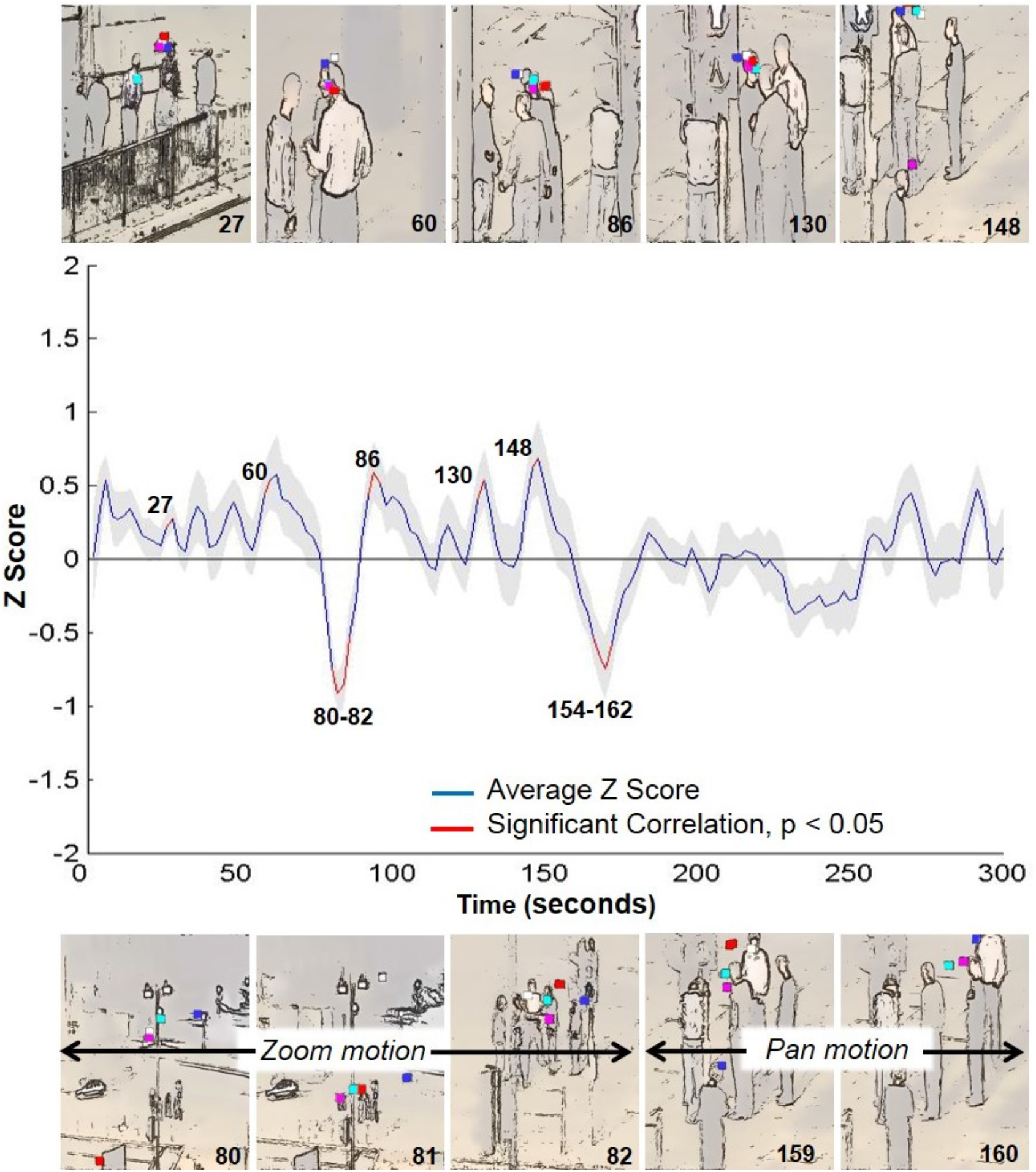
Reverse Correlation for FFA Activation in Experts for Prosocial Movie. The time course of FFA brain activity in expert viewers is used to illustrate the reverse correlation results with time frames indicated above each peaks and trough in brain activity significantly different from the mean value in red. Grey background denotes standard error around the mean activation. Participants’ eye movements are overlaid on the corresponding still images derived from the prosocial movie to illustrate the visual selectivity for faces (above) and the impact of motion on the visual availability of salient features.

**Fig 5.**
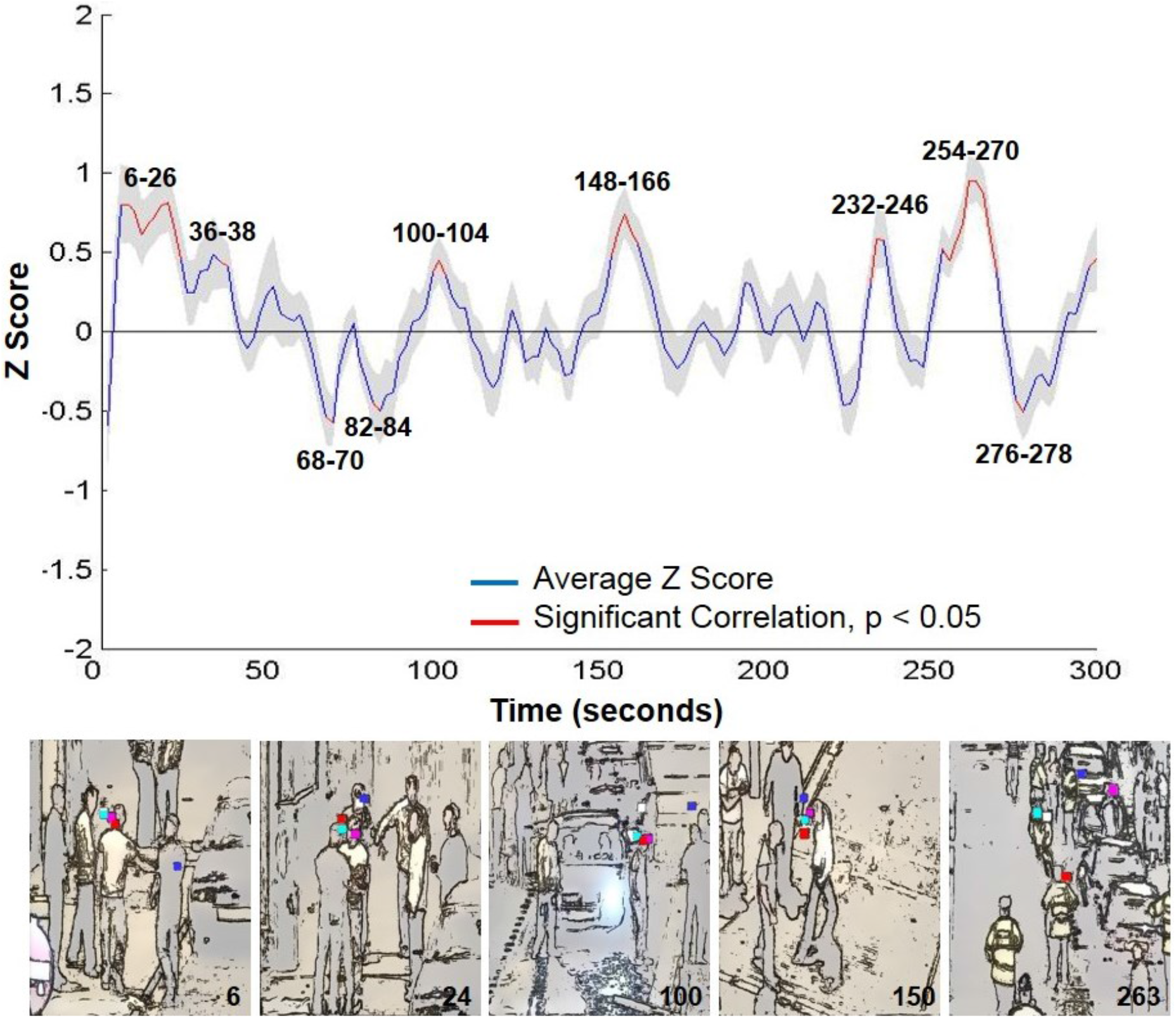
Reverse Correlation for FFA Activation in Experts for Antisocial Movie. The time course of FFA brain activity in expert viewers is used to illustrate the reverse correlation results with time frames indicated above each peaks and trough in brain activity significantly different from the mean value in red. Grey background denotes standard error around the mean activation. Participants’ eye movements are overlaid on the corresponding still images derived from the antisocial movie to illustrate the visual selectivity for faces (above).

#### Posterior Superior Temporal Sulcus (pSTS)

Reverse correlation revealed a strong association between increases in pSTS activity and the perception of intentional actions and body movements across the prosocial (Fig 6) and antisocial movie (Fig 7). Although the prosocial movie contained mainly commonplace and repetitive low-level social interactions, Fig 6 illustrates the overlap of eye fixations on salient body movements and the detrimental effect of large-scale camera movements. For example, one significant peak in pSTS activation was triggered by one individual unexpectedly throwing his hands up and down. In contrast, zooming out and camera tracking resulted in dips in activation.

**Fig 6.**
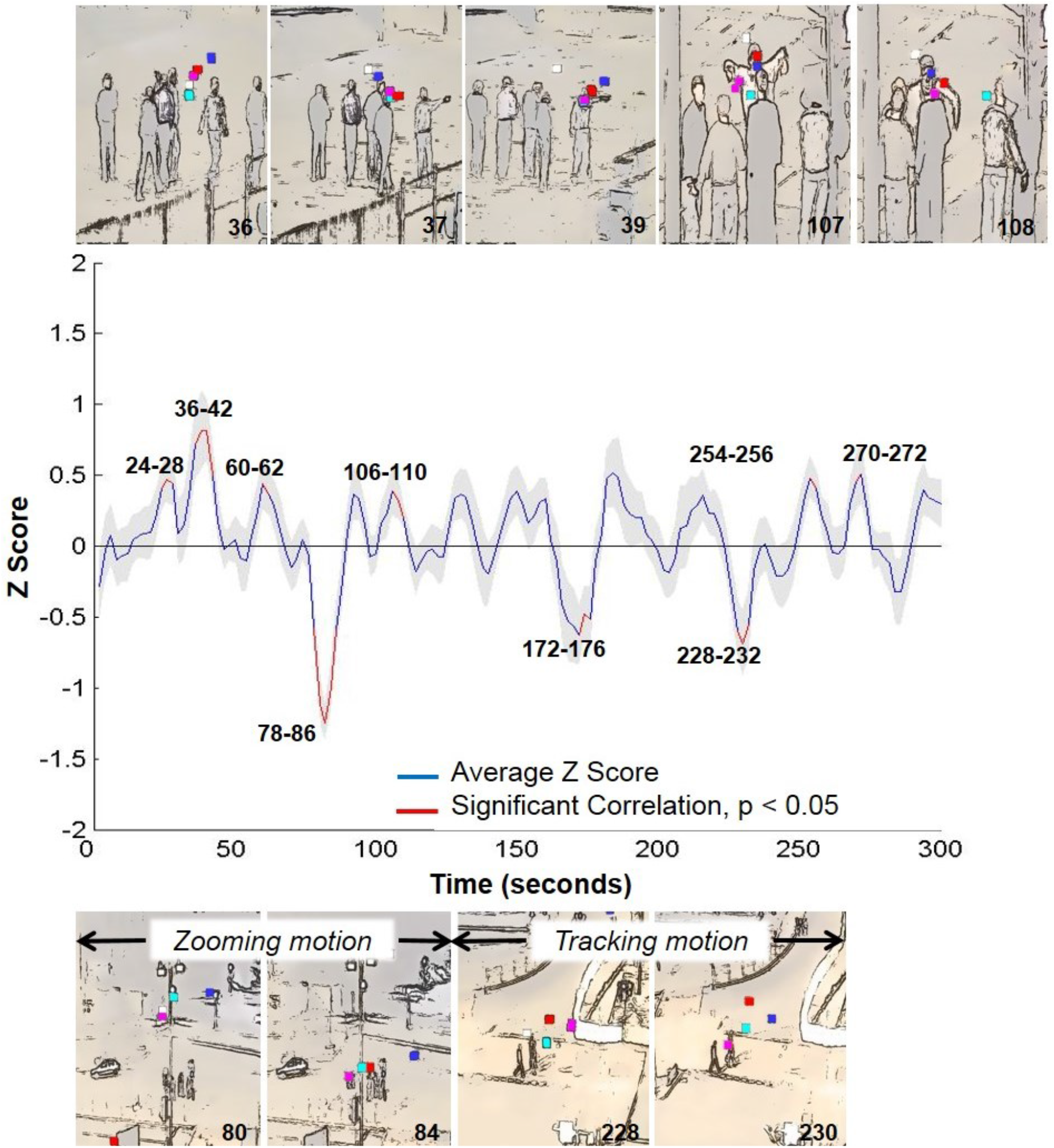
Reverse Correlation for pSTS Activation in Experts for Prosocial Movie. Reverse correlation results illustrated by eye movement data overlaid on still frames denoting significantly correlated time points. Still images above represent peaks in activation, while camera motion below, e.g. zooming and tracking correspond to troughs in activation. Grey background denotes standard error around the mean activation.

**Fig 7.**
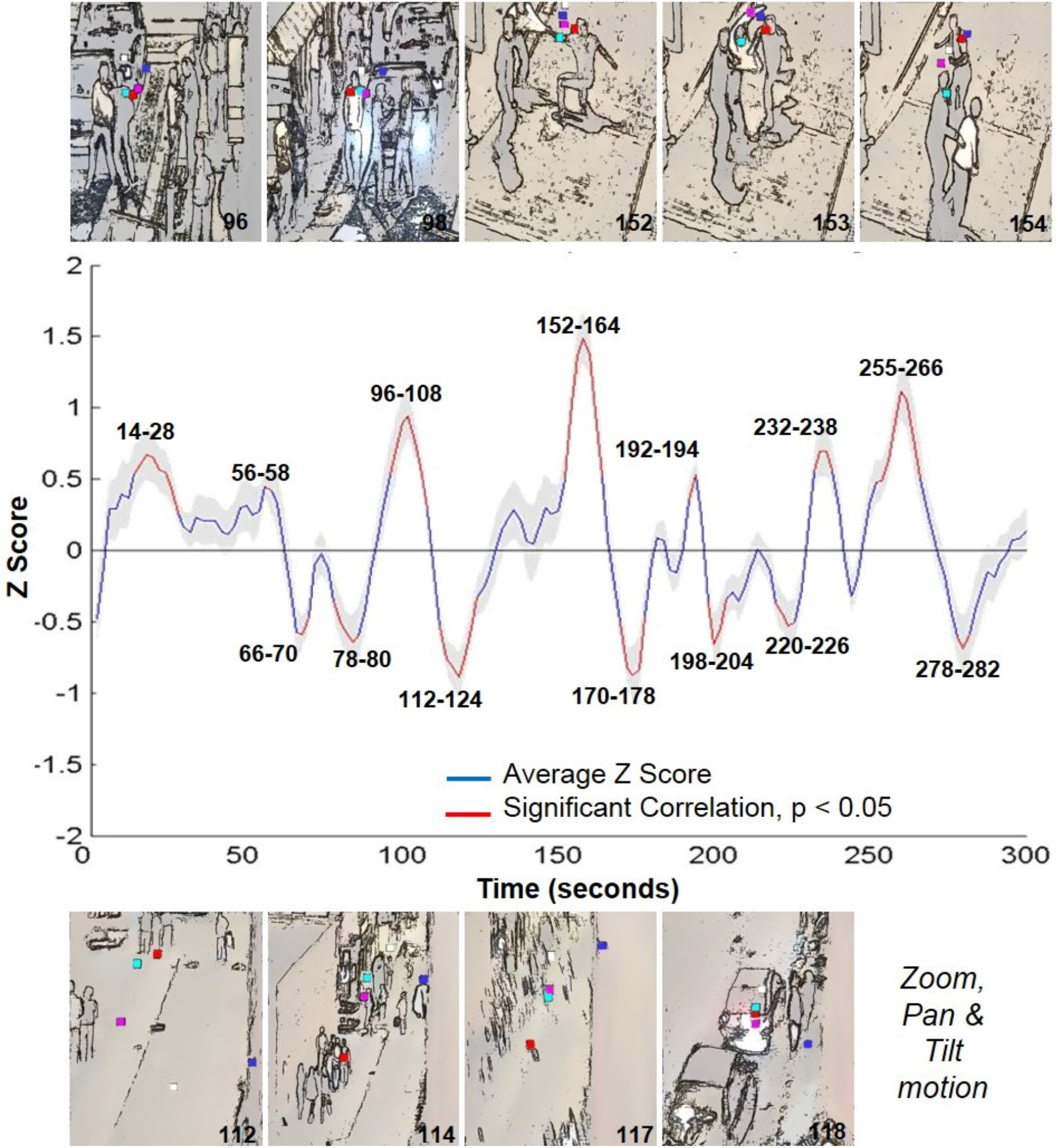
Reverse Correlation for pSTS Activation in Experts for Antisocial Movie. Reverse correlation results are illustrated by eye movement data overlaid on still frames denoting significantly correlated time points. Still images above represent peaks in activation for two instances of rapid antisocial actions representing a punching motion, while images below denote camera motion resulting in significant troughs in activation.

In contrast, the antisocial movie was characterized by a series of hostile interactions moving at a fast pace through an urban setting. This is reflected for instance in the largest significant peaks in pSTS activation observed in the antisocial movie. The viewers’ gaze closely follows the sequence of hand and body movements of one individual swinging a punch at his adversary, while in an earlier sequence two individuals are approaching each other aggressively in front of an on-coming vehicle. Similar to FFA activation, novices and experts show overlap in brain activity in response to viewing faces and/or body movements and are similarly affected by camera movements. However, in contrast to FFA activation, the camera motions elicited stronger significant dips in activation during the perception of non-biological motion patterns. These are in direct juxtaposition to significant peaks in activation during the perception of biological motion.

### Expert versus Novice Viewers

To further examine differences between viewer’s levels of expertise under naturalistic viewing conditions, the relative frequency of significantly activated time points in FFA and pSTS compared to the overall number of time points was used to investigate differences between expert and novice viewers while viewing antisocial and prosocial movies. In general, the antisocial movie was characterized by a higher percentage of significantly activated time points compared to the prosocial movie, in particular for pSTS activity. Moreover, novices tended to overlap more in significant peak activity as opposed to experts, in particular for the prosocial movie (Fig 8).

**Fig 8.**
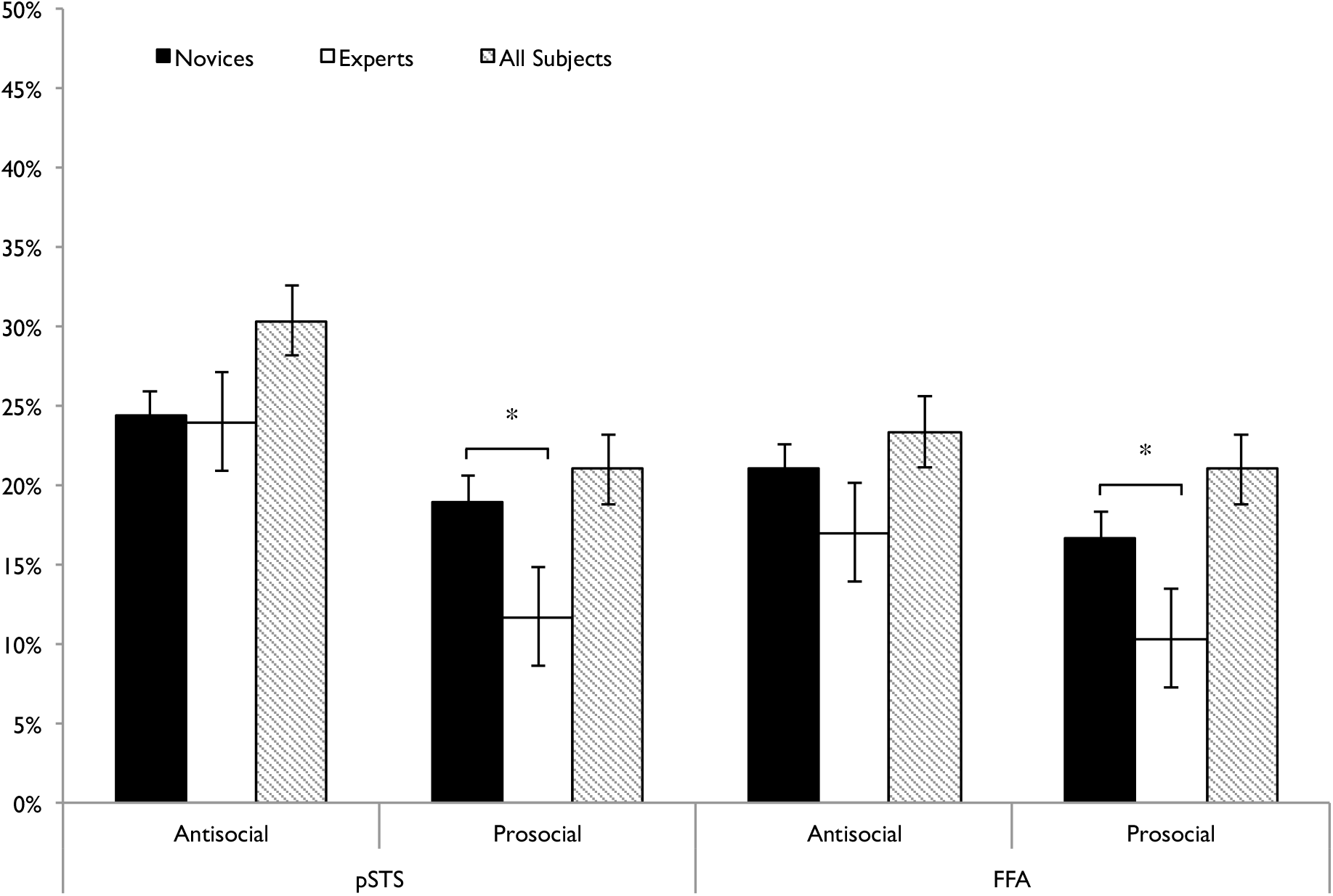
Percentage of Significantly Activated Timeframes. The percentage of significantly activated frames different from the average mean value (p<0.05) reflects an objective level of agreement between viewers. Error bars denote standard error. Novice viewers revealed a significantly higher percentage of intersubject agreement for the prosocial movie in both pSTS and FFA brain activity compared to experts.

Chi-squared analysis for significantly activations in FFA revealed a greater frequency in significantly activated timeframes in novice viewers compared to expert viewers in the prosocial movie (χ^2^ (2) = 8.413, p = 0.015), while viewing the antisocial movie revealed no significant relationship between significant FFA activation and viewer’s expertise (χ^2^ (2) =2.907, p = 0.234). Chi-squared analysis of the relationship between significant pSTS activity over time and viewers’ expertise revealed a similar pattern. There was a significant difference in frequency of significantly activated timeframes between viewers, with novice viewers activating more as compared to experts and all subjects combined viewing the prosocial movie (χ^2^ (2) =10.792, p = 0.005). In contrast, the antisocial movie revealed no significant relationship between viewers’ expertise and pSTS activity over time (χ^2^ (2) =3.011, p = 0.222).

## Discussion

In this study we used a reverse correlation analysis to investigate brain activity in novice and experienced CCTV observers when they viewed 300 second prosocial and antisocial videos of street activity. We focused on the FFA and pSTS regions as they are widely regarded to be involved in social processing and could be functionally localised. This neuroimaging analysis was supplemented by examination of the eye movements of a subset of the participants scanned. Our hypothesis stated that peak activation in FFA and pSTS would be significantly associated with the perception of human faces and human motion, while troughs in activation would be associated with large camera movements and the lack of visible faces or biological motion. Our findings, validated by the eye movement data, indicate that the predicted modulation of brain activity does occur as a result of salient features of faces and biological motion embedded within the naturalistic stimuli. The examination of expertise revealed that in both pSTS and FFA the novices had significantly more activated timeframes than the experienced observers for the prosocial video, which is in line with our prediction of reduced level of activation in CCTV operators. However, no difference was found for the antisocial video.

Our finding in FFA complements Hasson et al. (Hasson et al., 2004) who used reverse correlation to associate FFA activations with the presence of faces during the presentation of a cinematic movie. We were able to replicate the effect of reverse correlation in the context of face processing and increased FFA activation upon the perception of faces, even in qualitatively diminished stimuli, in which faces were not as readily available to visual perception as in Hasson’s cinematic displays of close-up shots. This finding serves to confirm our hypothesis that the eye movements we obtained centred on or around faces, and facial features correlated with peaks in localized brain activation within the FFA as revealed through the reverse correlation analysis. As with FFA, the peaks and troughs in pSTS activation similarly occur when salient features relating to human motion and movement were visible in the complex and dynamic CCTV footage. Although motion – of human and mechanical kind occurs almost constantly throughout the movies due to the nature of the camera technology, significant peak activity in pSTS corresponded specifically to the perception of biological motion, and in particular individual actions by the protagonists within the antisocial and prosocial movie as opposed to the perception of non-biological camera related motion.

Specifically, this study found that actions of intent that contained an affective component were more likely to trigger a significant peak in pSTS activation, in contrast to non-affective biological motion actions, such as walking down the street for instance. An example of affective biological motion was found in particular within the antisocial movie, in which an attacker is seen to prepare and execute a punch directed at his adversary. This sequence of events triggered an unusually large significant peak in pSTS activation, as it constitutes a movement with violent intent directed at another individual. This is in contrast to the remaining significant peaks, which focus on more typical biological motion, such as walking or running or simple group interactions, such as shaking hands, waving goodbye or jumping up and down, which were the majority of actions in the prosocial movie. Affective regulation of brain activity was also found in the study by Hasson et al. (Hasson et al., 2004) who argued that affective and surprising moments within the cinematic displays resulted in higher brain activity, as revealed by reverse correlation. Frith describes this as a mechanism determining the “social brain”, which is focused on biological agents as triggers for underlying mechanisms implemented in social interaction - of positive and negative nature [29].

In contrast to social cues from the face and body which resulted in peaks, large camera movements or camera devices, such as panning, zooming or cuts between cameras covering different viewpoints resulted in significant troughs in both FFA and pSTS activation. This result is consistent with the fact that salient features within the naturalistic scene became inaccessible or had moved out of the range of focus. It is also relevant to note that the occurrence of camera devices at times also corresponded to times with very little activity occurring within the scene, which in some instances may have contributed to the decrease in activation. However, recent results (Herbec et al., 2015) using unedited and edited videos of the same dance performance revealed that editing activates a wide network of brain areas including frontal cortex and possibly interactions within these brain networks might interact with pSTS and FFA activity.

Differences between the groups of observers were apparent in the results of the chi-squared analysis revealing a greater frequency of significantly activated timeframes for novices viewing the prosocial CCTV footage only compared to CCTV operators. This effect was true for both pSTS and FFA localizers, suggesting a similar demand on biological and face processing mechanisms throughout the movie. The nature of the prosocial movie depicted a small group of people centred within a large square in a city centre. Given a limited range of movements as a group throughout the majority of the footage, the visual availability of faces and biological motion were equally disrupted by large-scale cinematic devices, such as zooming out and panning. However, CCTV operators viewing the prosocial movies resulted in fewer significant peaks in FFA activation, suggesting a lesser degree of attentional synchronicity. In other words, it appears that novice viewers were more susceptible to focusing on the most common visual features, while expert viewers engage in a more in depth and complex visual analysis of facial and biological motion features across the scene, which is represented in the attentional variability. These results are similar to Petrini et al. (2014), who found increased activation in experienced relative to novice viewers for scenes depicting playful interactions. This result was consistent with informal reports of CCTV operators that they often attend closely to ambiguous playful interactions, which may degenerate into fights or violent behaviour.

In comparison, there was an equal level of synchronicity between observer groups in terms of pSTS activity or biological motion processing, which may be in part due to the fact that motion is perceived more as a whole and therefore can be affected by a range of motion rather than an activation which requires a specifically localised fixation on an individual face for instance. As there were multiple individuals displayed on the screen, the movement of the group as a whole will have affected both novice and expert viewers regardless of whom or which particular motion they were attending to. In addition, the antisocial movie was characterized by a faster pace and more aggressive nature potentially yielding a ceiling effect for the two groups, while in the slower paced prosocial video the efficiency indicated by experienced observers (Petrini et al., 2014) became apparent.

In summary, the research has shown that even in realistic natural scenes, the human brain can perceive and extract individual features from a complex dynamic environment. This is evident in the strong fMRI response recorded in areas specific to features such as face and biological motion processing. However, in contrast to previous research, this study has employed a data-driven approach to narrowing down the functionality of brain areas through the use of eye movements and correlated brain activity. Building on the literature for FFA activation the results have been extended to pSTS. These results indicate that motion, for instance camera movement, alone is insufficient in eliciting a strong fMRI result. Rather the presence of stimuli depicting biological motion and motion containing affective actions are crucial in eliciting a response with a special emphasis on social interactions. The results of our reverse correlation analysis advance the neuro-ergonomic approach of Grafton and Tipper (Grafton and Tipper, 2012) by showing how brain activity relates to complex real-world scenarios as viewed by novice and experienced viewers. The effect of CCTV experience may in addition result in more thorough dynamic scene processing compared to novice viewers for slower paced prosocial interactions, which may contain ambiguous social cues as compared to fast-paced explicitly hostile interactions that command visual attention equally across viewers.

This study was a novel approach in combining eye movement and reverse correlation using CCTV stimuli. There are, however, some limitations to the study. As with many such naturalistic stimuli it was impossible to precisely match the stimuli of the prosocial and antisocial stimuli. Although there were similarities in time of day, number and gender of those involved and camera movement, the antisocial video was faster paced and occurred in a more crowded environment. Furthermore, collecting eye movements in the MRI scanner is technically challenging and this precluded a complete set of eye movement data.

## Conclusion

In conclusion, we investigated the use of reverse correlation to indicate how brain activity in FFA and pSTS reflects activity in the viewing of naturalistic CCTV videos of prosocial and antisocial activity. We examined both novice and experienced CCTV observers. Reverse correlation results from FFA corroborated earlier results of Hasson and colleagues (Hasson et al., 2004) that used Hollywood style edited movies. As well, results from pSTS showed peaks related to intentional activity and troughs related to camera motion, underscoring the importance of processing dynamic social scenes. Finally, a difference was found between experienced and novice observers only on prosocial displays and this is consistent with previous findings about expertise (Petrini et al., 2014; Roffo et al., 2013) in that experience effects are reflected at higher levels of processing of dynamic naturalistic scenes.

## Acknowledgments

This work was funded by the Human Dimension and Medical Research Domain of the Dstl Programme Office for the UK Ministry of Defence. JG contribution was partially supported by the Sanford scholarship from the Department of Psychology, University of Glasgow. We thank all the CCTV staff and other participants for their contribution to this research.

## Data Statement

Due to the sensitive nature of the CCTV footage used in this study, raw data will remain confidential and cannot not be shared.

